# Calcineurin-fusion facilitates Cryo-EM Structure Determination of a Family A GPCR

**DOI:** 10.1101/2022.03.27.485993

**Authors:** Jun Xu, Geng Chen, Haoqing Wang, Sheng Cao, Jie Heng, Xavier Deupi, Yang Du, Brian K. Kobilka

**Affiliations:** Department of Molecular and Cellular Physiology, Stanford University School of Medicine, Stanford, CA 94305, USA; Kobilka Institute of Innovative Drug Discovery, School of Life and Health Sciences, Chinese University of Hong Kong, Shenzhen, Guangdong, 518172, China; Beijing Advanced Innovation Center for Structural Biology, School of Medicine, Tsinghua University, Beijing 100084, China; Condensed Matter Theory Group, Division of Scientific Computing, Theory, and Data, Paul Scherrer Institute, 5232 Villigen, Switzerland; Laboratory of Biomolecular Research, Division of Biology and Chemistry, Paul Scherrer Institute, 5232 Villigen, Switzerland; Swiss Institute of Bioinformatics (SIB), Lausanne, Switzerland

## Abstract

Advances in singe-particle cryo-electron microscopy (cryo-EM) have made possible to solve the structures of numerous Family A and Family B G protein coupled receptors (GPCRs) in complex with G proteins and arrestins, as well as several Family C GPCRs. Determination of these structures has been facilitated by the presence of large extra-membrane components (such as G protein, arrestin, or Venus flytrap domains) in these complexes that aid in particle alignment during processing of the cryo-EM data. In contrast, determination of the inactive state structure of Family A GPCRs is more challenging due to the relatively small size of the seven transmembrane domain (7TM) and to the surrounding detergent micelle that, in the absence of other features, make particle alignment impossible. Here we describe an alternative protein engineering strategy where the heterodimeric protein calcineurin is fused to a GPCR by three points of attachment, the cytoplasmic ends of TM5, TM6 and TM7. This three-point attachment provides a more rigid link with the GPCR transmembrane domain that facilitates particle alignment during data processing, allowing us to determine the structures of the β_2_ adrenergic receptor (β_2_AR) in the apo, antagonist-bound, and agonist-bound states. We expect that this fusion strategy may have broad application in cryo-EM structural determination of other Family A GPCRs.

## Main text

GPCRs have been challenging subjects for structural biology for a long time. First, with the exception of bovine rhodopsin, GPCRs are not sufficiently abundant in any mammalian tissue and have to be expressed in heterologous systems (*1, 2*). Also, most Family A GPCRs are dynamic flexible proteins with little exposed polar surface to facilitate the formation of a crystal lattice(*3, 4*). Furthermore, most GPCRs are relatively unstable, necessitating long-chain detergents with relatively large micelles (*3-5*) for efficient purification. Three strategies have been used to overcome these limitations: the development of antibodies that stabilize the GPCR and provide additional polar surfaces (*6*); thermostabilizing mutations that enable the use of short-chain detergents with smaller micelles (*7, 8*); and protein engineering to replace the flexible N-terminus or intracellular loop (ICL) 3 with highly crystallizable proteins such as T4 Lysozyme and BRIL (*9, 10*). Nevertheless, crystallization of Family A GPCRs still requires an element of luck and extensive rounds of optimization of the linkers between the receptor and the fusion protein, as well as optimization of crystallization conditions. Moreover, crystallization of GPCRs often depends on the availability of a high affinity ligand. In contrast, structure determination by cryo-EM is less dependent on luck. If the protein is of sufficient quality, stability, and size, a structure will likely be obtained. In this work, we sought to develop a protein engineering strategy that would enable the use of cryo-EM to determine structures of inactive-state Family A GPCRs that does not require the development of receptor-specific antibodies or nanobodies.

We chose to extend the ICL3 fusion protein strategy previously developed for crystallography (*9*) by adding an additional link through the C-terminus. We expected that the incorporation of a third link between the soluble protein and the receptor would reduce the flexibility between the two proteins and thereby improve its use as a fiducial marker for the 7TM core. This could be accomplished with a single protein having two or more independently folded domains or with two proteins that form a heterodimer (Fig. 1A). The proteins would have to have their N-and C-termini in positions that would be compatible with the relative positions of the cytoplasmic ends of TMs 5, 6 and 7 of the GPCR. Calcineurin (CN), composed of CN-A and CN-B subunits (Fig. 1B), was selected as our first candidate as it has the added advantage of being a Ca^2+^ dependent heterodimer (*11*). As shown in Fig. 1B, CN-B interacts with the C-terminal helix of CN-A (the CN-B binding region) in the presence of four Ca^2+^ ions. The link between CN-A and CN-B can be further stabilized by the FK binding protein (FKBP12) in the presence of the inhibitor FK506 (or Tacrolimus) (Fig. 1B).

**Fig. 1.**
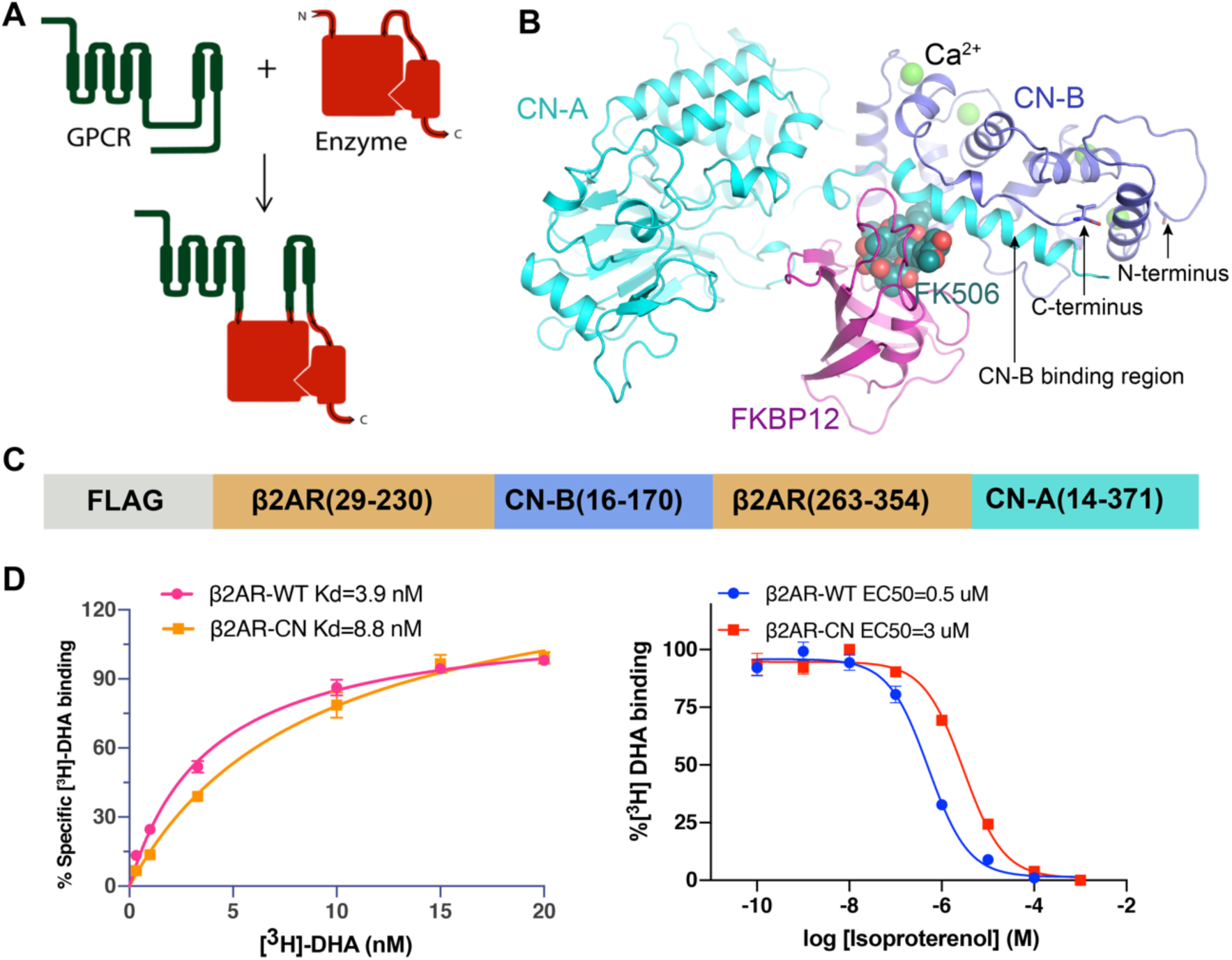
Engineering of the β2AR-CN fusion protein. (A) Concept design of the three-point fusion strategy. (B) Structure of the calcineurin heterodimer in complex with FKBP12 and FK506 (PDB: 1TCO). Ca^2+^ atoms are shown as green spheres. (C) Optimized construct of the β2AR-CN fusion protein. (D) Comparison of the ligand binding properties between the β2AR-CN-fusion construct and WT β2AR. Left panel: saturation binding; right panel: competition binding.

While the N and C-termini of CN-B are 19 Å apart in the structure of CN-FKBP12(*12*), the N-terminus is a 15 amino acid long flexible loop that could be conceivably shortened to accommodate the 11 Å distance between the cytoplasmic ends of TM5 and TM6 in the β_2_AR (Fig. 1B). Our initial construct is shown in Fig. S1A, where CN-B (amino acids 16-170) is inserted between amino acids 230-263 of the β_2_AR and the CN-A (amino acids 2-371) is fused to Y354 in the C-terminus of the β_2_AR. We were able to obtain a well-behaved monomeric fusion protein (Fig. S1B-C). 2D averages show clear architectures for β_2_AR, CN-A, and CN-B (Fig. S1D). 3D reconstruction finally yields a map with an overall resolution of 4.2 Å, which shows a relatively rigid orientation between the 7TM of β_2_AR and CN (Fig. S1E-G). Of note, computational docking of 3D models into the 4.2 Å map shows that the N-terminal ∼15 aa of CN-A lack observable density. This region was also not resolved in the crystal structure, pointing to an intrinsically disordered character (Fig. S1H). We further truncated the N-terminus of CN-A to create a shorter construct (Fig. 1C). Although density in β_2_AR is lost after position L341 (Fig. S1H), we kept the following residues (342-354) to better accommodate the distance between L341 and the N-terminus of CN-A. Radioligand binding studies show that the functional properties of the optimized construct are similar to those of wild-type β_2_AR, with almost the same binding affinity for the antagonist carazolol and moderately decreased binding affinity (6 fold) for the agonist isoproterenol (Fig. 1D).

To further stabilize the conformation of the CN heterodimer, we used the FKBP12 protein to form a complex with β_2_AR-CN (Fig. S2A). Fig. 2A shows a representative 2D class average of the complex, from which we can clearly see all the components of the fusion protein and the FKBP12. We finally reconstructed a 3.5 Å map from the dataset of the β_2_AR-CN-FKBP12 complex (Fig. S2B-F) that allowed us to build a model for most of the β_2_AR, CN-A, CN-B, and FKBP12 (Fig. 2B-C and S2G). Densities for the inverse-agonist carazolol and most residues in the orthosteric pocket are well resolved and the ligand can be docked with confidence (Fig. 2C-D). We also observe clear density for the compound FK506 which links FKBP12 with CN-A and CN-B (Fig. S2H). Moreover, the density for the linkers between CN-B and TM5 and TM6 are well-resolved (Fig. 2E). Although we didn’t observe a continuous density between the C-terminus of the β_2_AR and the N-terminus of CN-A in the final high-resolution map, the third linker was clearly seen in a well-defined low-resolution map from 3D classification (Fig. S2I).

**Fig. 2.**
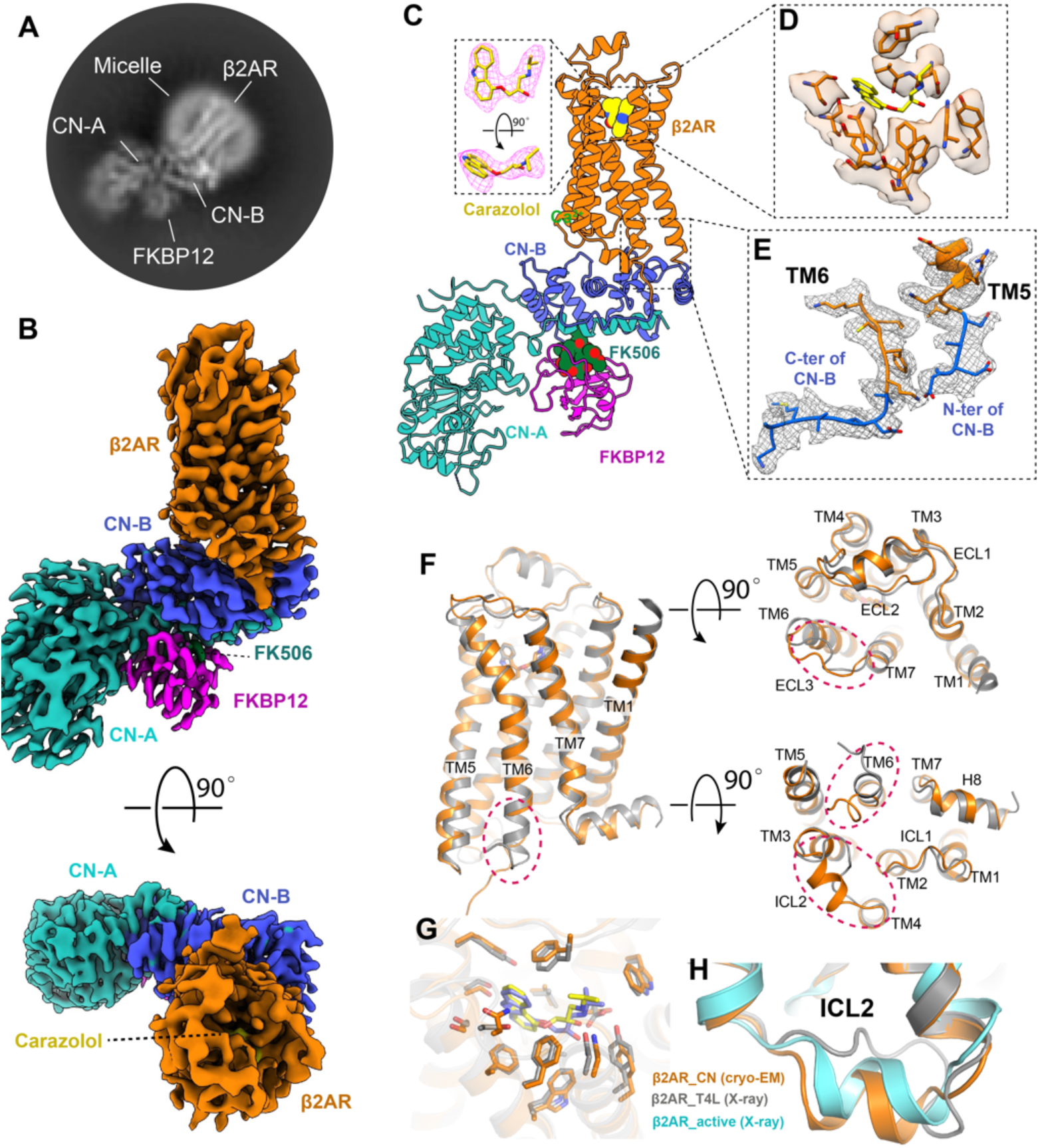
Cryo-EM structure determination of inverse agonist-bound β2AR using a three-point fusion strategy. (A) A representative 2D cryo-EM average of the β2AR–CN-FKBP12 complex shows high resolution features. (B) Cryo-EM density map of the β2AR–CN-FKBP12 complex from side view and top views. (C) 3D model of the β2AR–CN-FKBP12 complex. The inset shows the density corresponding to the inverse agonist carazolol depicted as magenta mesh. The map and model in all panels are colored according to polypeptide chains. (D-E) Density maps and models of the residues in the orthosteric binding pocket (D) and the TM5/TM6 linkers between β2AR and CN-B (E). (F) Overall structural comparison between the cryo-EM structure (orange) and X-ray crystal structure (gray) of the β_2_AR. Differences are highlighted with red circles. (G) Comparison of the carazolol binding pocket between the cryo-EM and crystal structures. (H) Comparison of the ICL2 conformation between inactive and active structures.

We next compared the cryo-EM structure of carazolol-bound β_2_AR with the 2.4 Å crystal structure of the β_2_AR-T4L fusion protein (PDB: 2RH1). As expected, the overall structures are similar, with a root mean square deviation (RMSD) of 0.7 Å (Fig. 2F). The carazolol binding pockets for the two structures are also highly similar, with a RMSD of 0.5 Å (Fig. 2G). The largest difference between the two structures corresponds to intracellular loop 2 (ICL2), which is folded as a helix in the β_2_AR-CN cryo-EM structure while it is an unstructured loop in the β_2_AR-T4L crystal structure (Fig. 2F and 2H). Existing structures of β_2_AR show that ICL2 is a helix in the active state (Fig. 2H) and that the transition from a loop to a helix conformation may be an important step for receptor activation (*13, 14*). Our result suggests that the helical conformation of ICL2 can also exist in the inactive state of β_2_AR, possibly in equilibrium with an unstructured loop conformation. The complete absence of the helical conformation of ICL2 in crystal structures of the inactive β_2_AR might be due to crystal lattice contacts with T4L (Fig. S3). Similarly, the slight difference in the conformation of ECL3 may also result from crystal packing (Fig. S3). In addition, we also observed different conformations at the cytoplasmic end of TM6, which is likely caused by the different fusion proteins linked to this helix (Fig. 2F).

Although crystallization has enabled the determination of many inactive GPCR structures, most of the successful cases required a high-affinity antagonist or extensive thermostabilization by mutagenesis to make the receptor as stable as possible(*3*). For most receptors –including the β_2_AR– it has not been possible to obtain structures of the apo or agonist-bound receptor alone by crystallography (in the absence of thermostabilizing mutations) due to the inherent dynamics of these states(*15*). Using the CN-fusion strategy, we were able to obtain the cryo-EM map of the β_2_AR in its apo-form and norepinephrine-bound form at 3.9 Å and 3.6 Å, respectively (Fig. 3A-B, S4 and S5). Due to the lower resolution of the apo-state, especially in the extracellular ends of the TM segments and extracellular loops (Fig. S4D), many of the side chains could not be modeled; nevertheless, the overall structures of the apo-state and norepinephrine-bound β_2_AR are nearly identical to the antagonist-bound structure (Fig. 3C). However, local resolution analysis show that the apo-state receptor is much more flexible than the antagonist-bound or agonist-bound receptor, especially in the orthosteric pocket and extracellular loops (Fig. 3D), consistent with the notion that ligand binding stabilizes the conformation around the orthosteric binding pocket. Of note, the TM6 conformation in the norepinephrine-bound structure is the same as the inactive state (Fig. 3C). Previously we obtained a crystal structure of the β_2_AR-T4L fusion protein bound to a covalent agonist where TM6 was also in an inactive conformation(*16*). These results are consistent with fluorescence and spectroscopic studies demonstrating that agonist alone cannot stabilize TM6 of the β_2_AR in a fully active conformation (*15, 17*). It’s also possible that the CN-fusion stabilizes the inactive conformation of TM6.

**Fig. 3.**
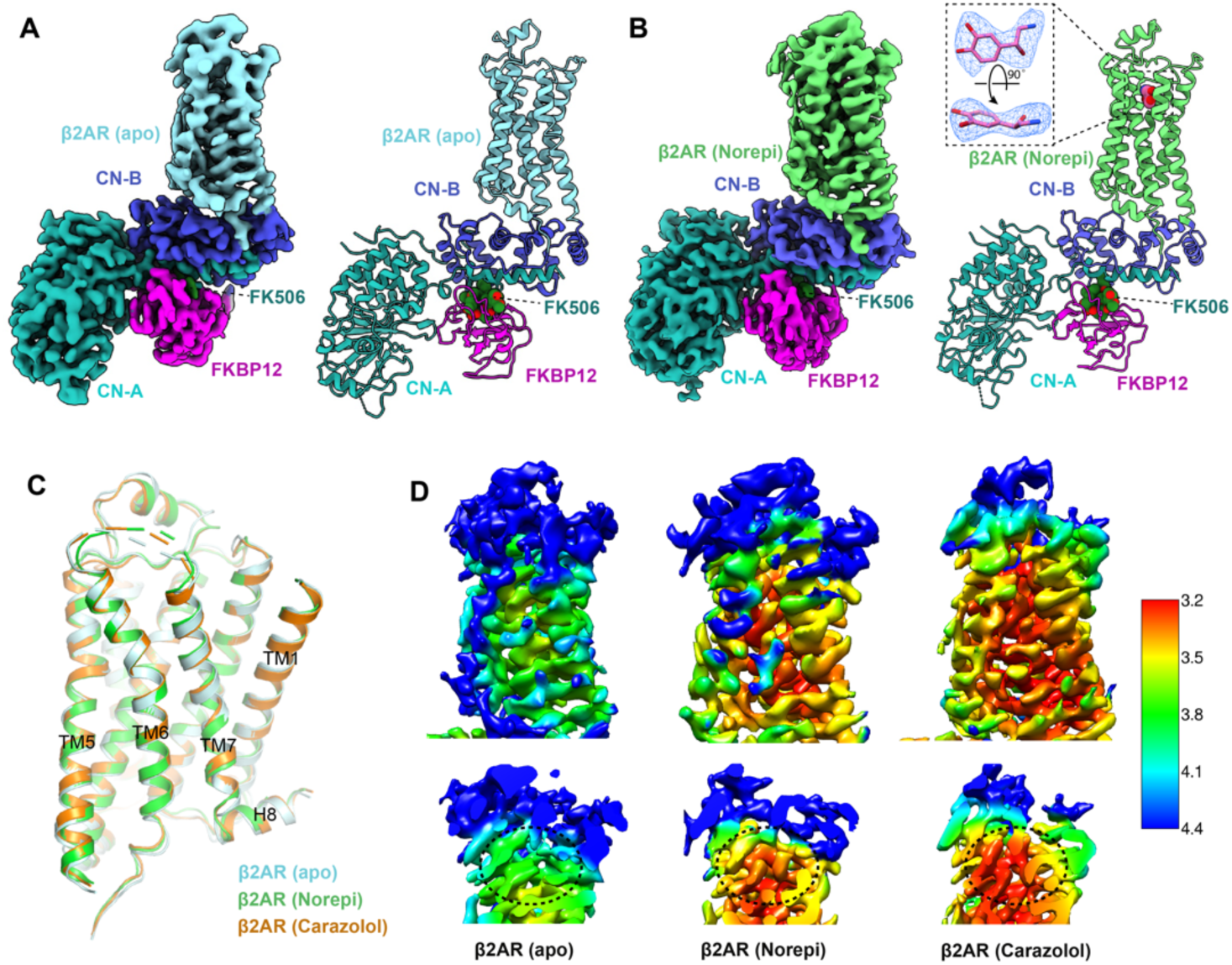
Cryo-EM structure determination of apo-state and agonist-bound β2AR using the CN fusion strategy. (A-B) Three-dimensional maps and models of the β2AR–CN-FKBP12 complex in apo (A) and norepinephrine-bound (B) states. The density map for norepinephrine is shown in the inset as a blue mesh. (C) Superimposition of the β2AR structures in apo (cyan), norepinephrine-bound (green) and carazolol-bound (orange) states. (D) Comparison of the local resolution maps of the β2AR in apo, norepinephrine-bound, and carazolol-bound states. The orthosteric binding pocket is highlighted with black circles in the bottom panels.

Pharmacological studies have revealed that agonists of GPCRs have at least two binding modes: a low affinity binding mode in the absence of intracellular proteins (e.g. G protein, arrestin, or nanobody) and a high affinity binding mode when coupled to intracellular binders(*18, 19*). This can be explained by the model of allosteric coupling between the intracellular signaling proteins and the ligand-binding pocket(*18*). Indeed, numerous biophysical studies have revealed the conformational changes stabilized by the intracellular protein propagate to the extracellular ligand binding site of GPCRs(*20, 21*). The norepinephrine-bound inactive β_2_AR structure, together with the previously determined active β_2_AR crystal structure bound to epinephrine –a nearly identical ligand– and stabilized by a G protein mimetic nanobody(*14*) allow us to compare the two states (low affinity and high affinity). For the norepinephrine-bound structure, densities for the ligand and orthosteric pocket residues are relatively well-defined (Fig. 3B and 4A). Comparison of the inactive and active structures shows that there is a 1.5-2 Å inward movement for TM5 and TM6 in the pocket upon binding with a G protein mimetic nanobody (Fig. 4B). The contraction of TM5 and TM6 results in different polar interactions between the ligand and receptor (Fig. 4C-D). The amino-ethanol group of norepinephrine/epinephrine forms similar hydrogen-bonding interactions with D113^3.32^ and N312^7.39^ in both states (Fig. 4C-D). In the active state, the meta-hydroxyl can form hydrogen bonds with both S203^5.42^ and N293^6.55^ while the para-hydroxyl forms hydrogen bond with S207^5.46^. S204^5.43^ is also involved in the polar network through a hydrogen-bonding interaction with N293^6.55^, which further stabilizes the interactions between TM5/6 and the ligand (Fig. 4D). In contrast, in the inactive state, norepinephrine can only form a weak hydrogen bond (∼3.5 Å) with S203^5.42^ through the para-hydroxyl, which results in much weakened overall interactions between receptor and the ligand (Fig. 4C). Together, these results provide a direct structural explanation for the distinct binding affinity of an agonist in the absence and presence of G protein or G protein mimetic nanobody.

**Fig. 4.**
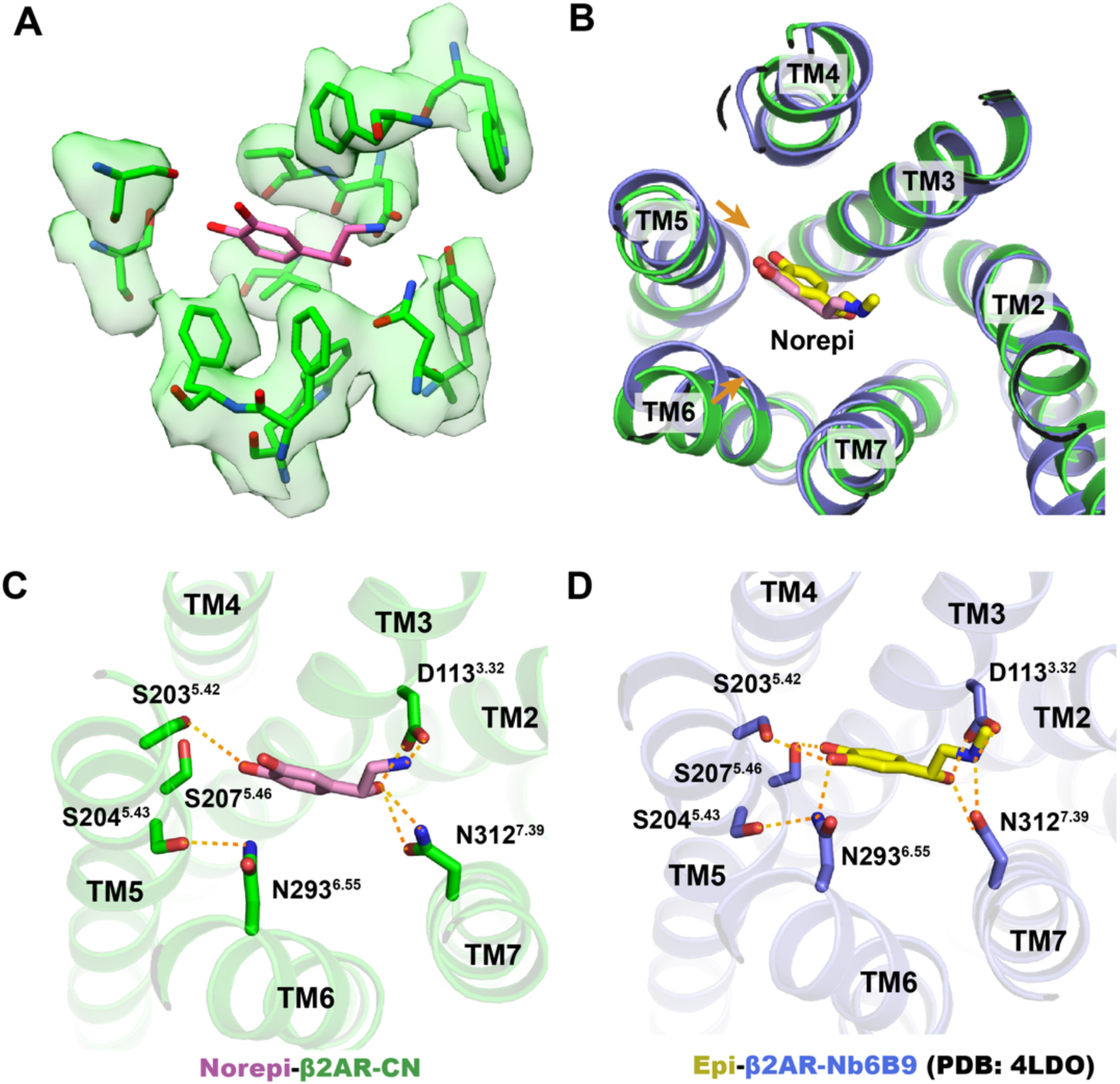
Comparison of agonist binding modes in inactive and active states. (A) Density map and model of the norepinephrine-bound orthosteric pocket residues. (B) Comparison of the orthosteric binding pockets of norepinephrine-bound inactive structure and epinephrine-bound active structure. (C-D) Detailed polar interactions between norepinephrine with inactive β2AR and epinephrine with active β2AR.

In summary, we have developed a protein engineering strategy for structural determination of GPCRs using cryo-EM, in which the heterodimer protein calcineurin is fused to a GPCR by three points of attachment. We demonstrated the feasibility of this method by solving the antagonist-bound inactive structure of the β_2_AR. Moreover, we show that this fusion strategy can also be utilized to determine the structure of receptors in the apo-state and bound to a low affinity agonist. Recently, a three-points fusion strategy has also been used to determine the high-resolution structure of a glucose transporter by fusion a GFP to the intracellular loop and a GFP-binding nanobody to the C-terminus(*22*). Although this particular study uses two different proteins interacting with each other, these results, together with our calcineurin-fusion strategy, suggest that the three-points fusion strategy might provide a more rigid fiducial maker for particle alignment of small integral membrane proteins during cryo-EM data processing. However, one might still need to optimize the three linkers to get the fusion protein as rigid as possible without affecting the receptor activity. While this might be a time-consuming step, the β_2_AR-CN design can be used as an initial template for most of other Family A GPCRs through simple sequence alignment. It should be noted that much higher resolution GPCR maps were obtained in a recent published study where a universal nanobody was used as a fiducial maker for cryo-EM analysis(*23*). This is probably due to the more rigid nanobody-bound receptor complex. Indeed, the local resolution map of β_2_AR-CN shows much better resolution of the CN than of the β_2_AR (Fig. S2F, S4D and S5D), indicating there is still flexibility between CN and receptor. Additional optimization of the linkers maybe helpful to further improve the resolution. We believe this approach may have broad applications in determining the structures of other Family A GPCRs in apo-state or ligand-bound (orthosteric and/or allosteric) state, and to facilitate structure-based drug development targeting GPCRs.

## Author Contributions

J.X., G.C., H.W., S.C. and J.H. performed experiments. X. D., Y.D. and B.K.K. conceived the project. Y.D. and B.K.K. provide overall supervision of the project. B.K.K. and J.X. wrote the manuscript with contributions from all authors.

## Acknowledgement

We would like to thank the Cryo-EM Center of the Chinese University of Hong Kong, Shenzhen for data collection. Some CryoEM data collection was performed at the Stanford-SLAC Cryo-EM Facilities, supported by Stanford University, SLAC, and the National Institutes of Health S10 Instrumentation Programs. The authors thank Elizabeth Montabana, Chensong Zhang and Dr. Zheng Liu for their support on CryoEM data collection. Part of CryoEM data processing for this project was performed on the Sherlock cluster. We would like to thank Stanford University and the Stanford Research Computing Center for providing computational resources and support that contributed to these research results. Y.D. is supported by grants from Science, Technology and Innovation Commission of Shenzhen Municipality (Project code 10120210014), and in part by the startup fund of Institute of Innovative Drug Discovery, Chinese University of Hong Kong, Shenzhen. This work was also supported by the American Heart Association Postdoctoral Fellowship (H.W.). B.K.K. is a Chan Zuckerberg Biohub Investigator.

## Materials availability

Plasmids are available from the corresponding authors upon request.

## Methods

### Expression and purification of FKBP12

The human FKBP12 gene was cloned into pGEX-2TK vector with a C-terminal 6x His tag. Plasmids were transformed into *E. coli* BL21(DE3) cells. Cells were grown to A600 = 0.8 at 37°C in TB media containing 0.1% glucose, 2 mM MgCl_2_ and 50 mg/ml ampicillin. Cells were induced by addition of 0.5 mM IPTG and were incubated for 8 hours at 37°C. Cells were harvest and disrupted by sonication. FKBP12 proteins were first purified by Ni-NTA chromatography and then size exclusion chromatography (SEC) with a buffer containing 20 mM HEPEs pH 7.5, 100 mM NaCl, 100 μM TCEP and 5 μM FK506. Monodisperse peak fractions were collected and concentration using a 3 kDa molecular weight cutoff Millipore concentrator to a concentration at around 15mg/ml. The concentrated FKBP12 were aliquoted, flash frozen in liquid nitrogen and stored at -80 °C before use.

### Expression and purification of β2AR-CN

The β2AR-CN fusion sequence was cloned into pFastBac vector with a N-terminal FLAG tag. Recombinant baculovirus for insect cell expression was made using the Bac-to-Bac system. Sf9 cells were grown in SIM SF Medium (Sino Biological Inc.) at 27 °C and were infected with recombinant baculovirus containing β2AR-CN gene at a density of 4 × 10^6^ cells per mL in the presence of 2 μM alprenolol. After 48 hours infection, the cells were spun down and cell pellets were stored at -80 °C until use. Thawed cell pellets were resuspended in a lysis buffer composed of 10 mM Tris, 1 mM EDTA, 10 μM carazolol or 100 μM norepinephrine, 2.5 μg/mL leupeptin and 160 μg/mL benzamidine to lyse the cells by hypotonic. Cell membranes were then spun down and solubilized with a buffer of 20 mM HEPEs pH 7.5, 100 mM NaCl, 1 % DDM, 0.03 % CHS, 2.5 μg/mL leupeptin, 160 μg/mL benzamidine, 5 mM CaCl_2_ and 10 μM carazolol or 100 μM norepinephrine at 4 °C for 1 hour. The solubilized receptor was then loaded onto a column with anti-flag M1 affinity resin and was extensively washed with a buffer containing 20 mM HEPEs, pH 7.5, 300 mM NaCl, 0.1 % DDM, 0.003 % CHS, 2 mM CaCl_2_ and 10 μM carazolol or 100 μM norepinephrine. The receptor was then gradually exchanged into a buffer containing 20 mM HEPEs pH 7.5, 100 mM NaCl, 0.0075% lauryl maltose neopentyl glycol (LMNG, Anatrace), 0.0025%GDN, 0.001% CHS, 2 mM CaCl_2_ and 10 μM carazolol or 100 μM norepinephrine, and then eluted with same buffer supplemented with 0.2 mg/ml flag peptide and 5 mM EDTA. The flag affinity chromatography purified receptor was supplemented with 10 mM CaCl_2_ immediately, then concentrated to 500 μL and finally purified by SEC chromatography with a buffer containing 20 mM HEPEs pH 7.5, 100 mM NaCl, 0.00075%LMNG, 0.00025%GDN, 0.0001% CHS, 0.5mM CaCl_2_ and 10 μM carazolol or 100 μM norepinephrine. The monodisperse peak fractions was collected and concentrated to ∼10 mg/ml for cryo-EM analysis. For complexing with FKBP12, the monodisperse peak fractions were collected and incubated with excess FKBP12 in the presence of 5 μM FK506 for 1 hour on ice. The β2AR-CN-FKBP12 mix was then subjected to SEC chromatography against a buffer containing 20 mM HEPEs pH 7.5, 100 mM NaCl, 0.00075%LMNG, 0.00025%GDN, 0.0001% CHS, 0.5mM CaCl_2_, 10 μM carazolol or 100 μM norepinephrine and 5 μM FK506. The complex peak was collected and concentrated to ∼10 mg/ml for cryo-EM. For the apo-state sample, no ligand was added in expression and all purification steps.

### Cryo-EM sample preparation and data collection

For carazolol and norepinephrine-bound samples, amorphous alloy film (CryoMatrix nickel titanium alloy film, R1.2/1.3, Zhenjiang Lehua Electronic Technology Co., Ltd.) was glow-discharged for 60 s at a Tergeo-EM plasma cleaner. 3 μL purified β_2_AR-CN or β_2_AR-CN-FKBP12 sample was applied onto the grid and then blotted for 3 s with blotting force 0 and quickly plunged into liquid ethane cooled by liquid nitrogen using Vitrobot Mark IV (Thermo Fisher Scientific, USA) at 4°C and with 100% humidity. Cryo-EM data were collected on a 300 kV Titan Krios Gi3 microscope. The raw movies were recorded by Gatan K3 BioQuantum Camera at a magnification of 105 000, and a pixel size of 0.85 Å. Inelastically scattered electrons were excluded by a GIF Quantum energy filter (Gatan, USA) using a slit width of 20 eV. The movie stacks were acquired with a defocus range of -1.0 to -1.6 micron with a total exposure time 2.5s fragmented into 50 frames (0.05s/frame) and with a dose rate of 22.0 e/pixel/s. The imaging mode was super resolution with 2-time hardware binning. Semi-automatic data acquisition was performed using SerialEM.

For the apo-state sample, a Quantifoil grid (R1.2/1.3, Au) was glow-discharged for 45 s at easiGlow discharged cleaning system. An aliquot of 3 μL sample was deposited onto the grid and plunge-frozen into liquid ethane using an Vitrobot Mark IV. Data collection was conducted on Titan Krios operated at 300 keV using a nominal magnification of 130,000x. Movies were captured using a Gatan K3 Summit direct electron detector in counted mode, which resulted in a pixel size of 0.85 Å. Movie stacks were obtained with a defocus range of -1.0 to -2.0 μm, using SerialEM 3.7.10 with a set of customized scripts enabling automated low-dose image acquisition. Each movie stack was recorded for a total of 8 seconds with 0.2 s per frame. The exposure rate was 7 electrons per pixel per second.

### Cryo-EM image analysis and model building

The image stacks of the β_2_AR-CN fusion protein were collected and subjected for motion correction using MotionCor2(*24*). Contrast transfer function parameters were estimated by CTFFIND4(*25*), implemented in RELION3.1(*26*). 2,000 particles were manually picked and extracted from the motion-corrected micrographs followed with 2D classification. Templates were selected from the 2D classification result. Particles were auto-picked using the templates in RELION and then subjected to 2D classification using cryoSPARC(*27*). Selected particles with appropriate 2D average from 2D classification were further subjected to Ab-initio reconstruction. Particles with appropriate initial model were selected from Ab-initio followed by homogeneous refinement in cryoSPARC. After global and local CTF refinement, the particles (kept to 325’064) were subjected to non-uniform refinement for a 4.20 angstrom reconstruction determined by gold standard Fourier shell correlation using the 0.143 criterion. Data processing for the β_2_AR-CN-FKBP12 complexes were all done in cryoSPARC. For the carazolol-bound β_2_AR-CN-FKBP12 complex, a total of 6’001 image stacks were collected and subjected to patch motion correction and patch CTF refinement. 2’819’890 particles were auto-picked using the β_2_AR-CN map as a template and then subjected to 2D classification followed by Ab-initio reconstruction and 2 rounds of heterogeneous refinement. The resulting particles were subject to non-uniform refinement and local refinement and yielded a map at 3.6 Å. After local motion correction and another round of non-uniform refinement and local refinement, the resolution was improved to 3.49 Å. The norepinephrine-bound and apo-state datasets were processed in a similar way as the carazolol-bound dataset(Fig. S4B and S5B). The crystal structure of the inactive β_2_AR (PDB code: 2RH1), rat calcineurin (PDB code: 4IL1) and FKBP12-FK506 (PDB code: 1FKJ) were used as initial models for model rebuilding and refinement against EM density map. The models were docked into the EM density map using UCSF Chimera(*28*), followed by iterative manual building in Coot(*29*) and refinement in Phenix(*30*).

### Radio-ligand binding assay

Cell membranes were prepared following a previously reported protocol(*31*). Radioligand saturation binding of the antagonist [^3^H]-dihydroalprenolol (PerkinElmer) was conducted with the cell membranes to a final volume of 250 μL. Briefly, the cell membranes were diluted in assay buffer containing 20 mM Hepes, pH 7.4, 100 mM NaCl, and 0.5 mg/mL BSA. Serial dilutions of [^3^H]-dihydroalprenolol at indicated final concentrations were incubated with the membranes for 1.5 hours while shaking at 200 rpm at room temperature. The membranes were washed and collected by filtration with a 48-well Brandel harvester. The filter papers containing the membrane were incubated with a 3 mL OptiPhase HiSafe 3 liquid scintillation cocktail. Radioactivity was counted with a Microbeta2 scintillation counter. All curves were constructed from mean ± s.e.m of three independent experiments at three replicas, and the saturation binding data were analyzed in the GraphPad Prism software using a one-site saturation binding equation. For radioligand competition binding, membrane solutions were incubated with 2nM [^3^H]-dihydroalprenolol and the unlabeled competitor to a final volume of 250 μL. The assay buffer contains 20 mM Hepes, pH 7.4, 100 mM NaCl, and 0.5 mg/mL BSA. Binding reactions were incubated and shaken at room temperature for 1.5 hours and were harvested using a 48-well Brandel harvester. All curves were constructed from mean ± s.e.m of three independent experiments at three replicas. Competition binding data were analyzed in the GraphPad Prism software using a one-site competitive binding equation.

## Supplementary Figures

**Fig. S1.**
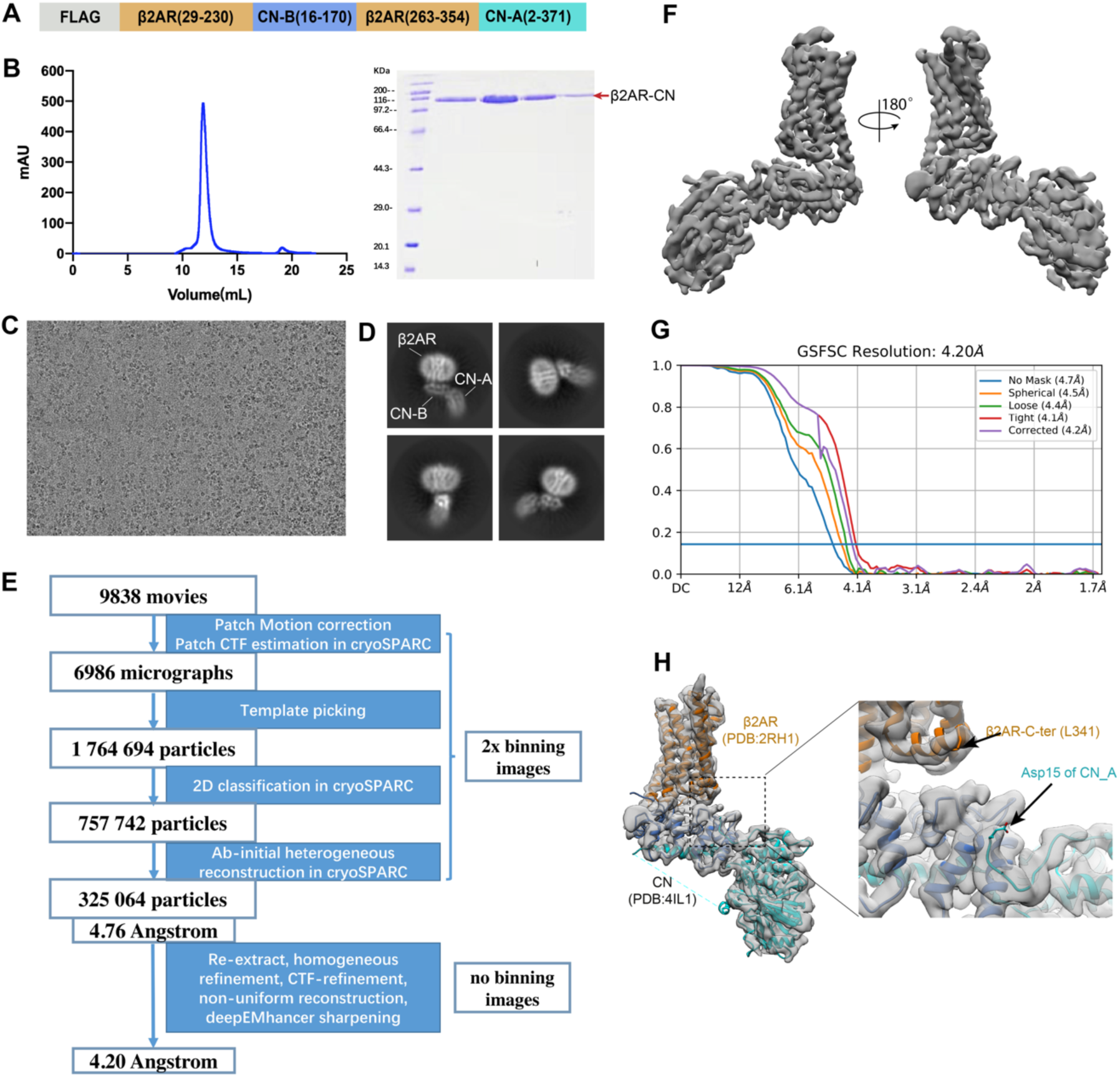
Sample preparation and cryo-EM data processing of the β2AR-CN fusion protein. (A) Design of the β2AR-CN construct. (B) Size exclusion chromatography profile and SDS-PAGE of the β2AR-CN fusion protein. (C) Representative micrograph of the β2AR-CN particles. (D) Representative 2D classification result. (E) Cryo-EM image processing workflow for image processing. (F) 3D cryo-EM map of the β2AR-CN fusion protein viewed from two directions. (G) Fourier shell correlation (FSC) curve with the estimated resolution according to the gold standard. (H) Docking of inactive β2AR (pdb:2RH1) and CN (pdb: 4IL1) into the cryo-EM density map.

**Fig. S2.**
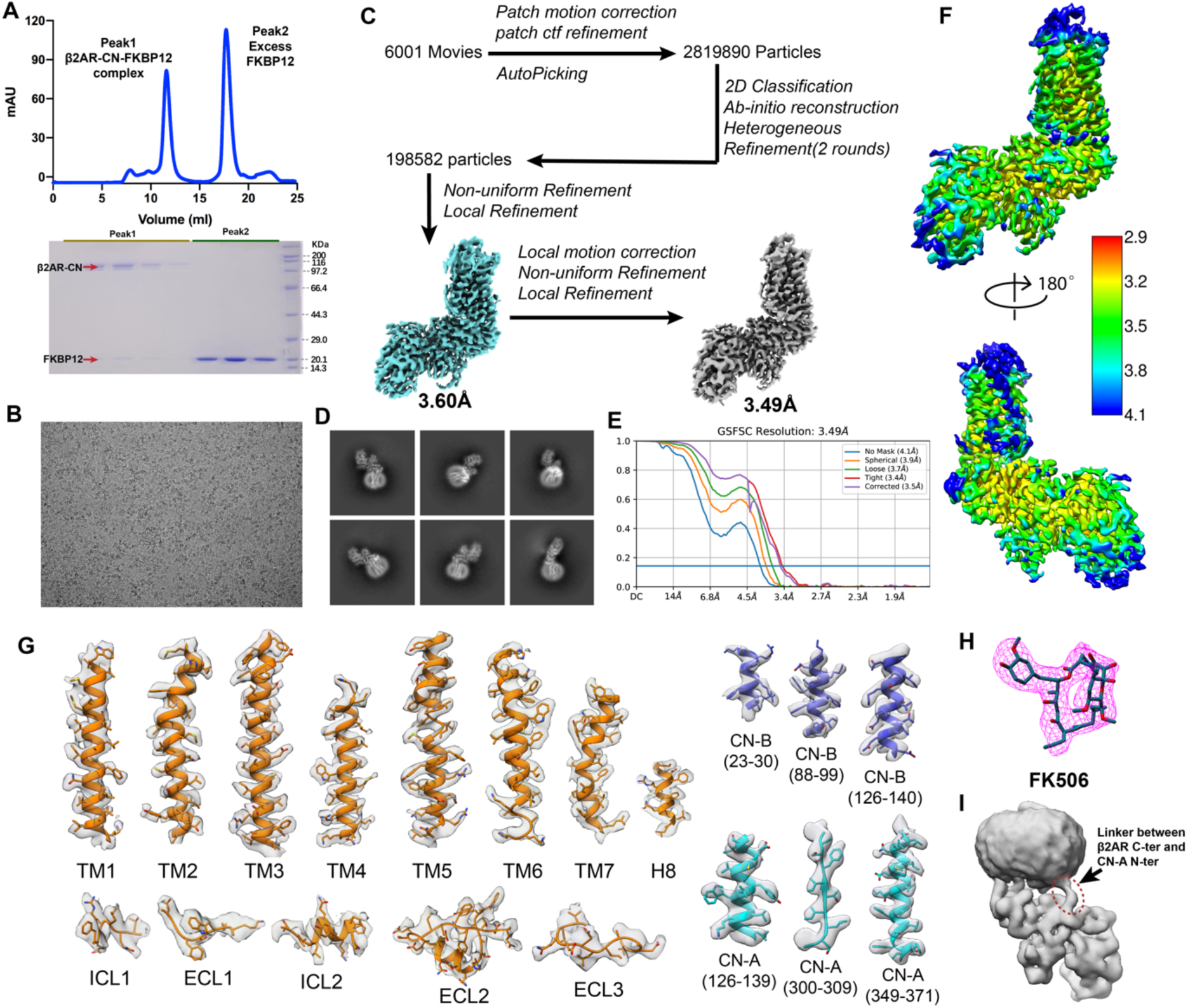
Sample preparation and cryo-EM data processing of carazolol-bound β2AR-CN-FKBP12 complex. (A) Size exclusion chromatography profile and SDS-PAGE of the complex. (B) Representative micrograph of the β2AR-CN particles. (C) Cryo-EM image processing workflow for image processing. (D) Representative 2D classification result. (E) Fourier shell correlation (FSC) curve with the estimated resolution according to the gold standard. (F) Local resolution map viewed from two directions. (G) Representative density maps and models for TM1-7, H8 and loop regions for β2AR and selected regions for CN. (H) Density map for FK506. (I) The linker between the β2AR C-terminus and CN-A N-terminus is shown in a well-defined low-resolution 3D classification map.

**Fig. S3.**
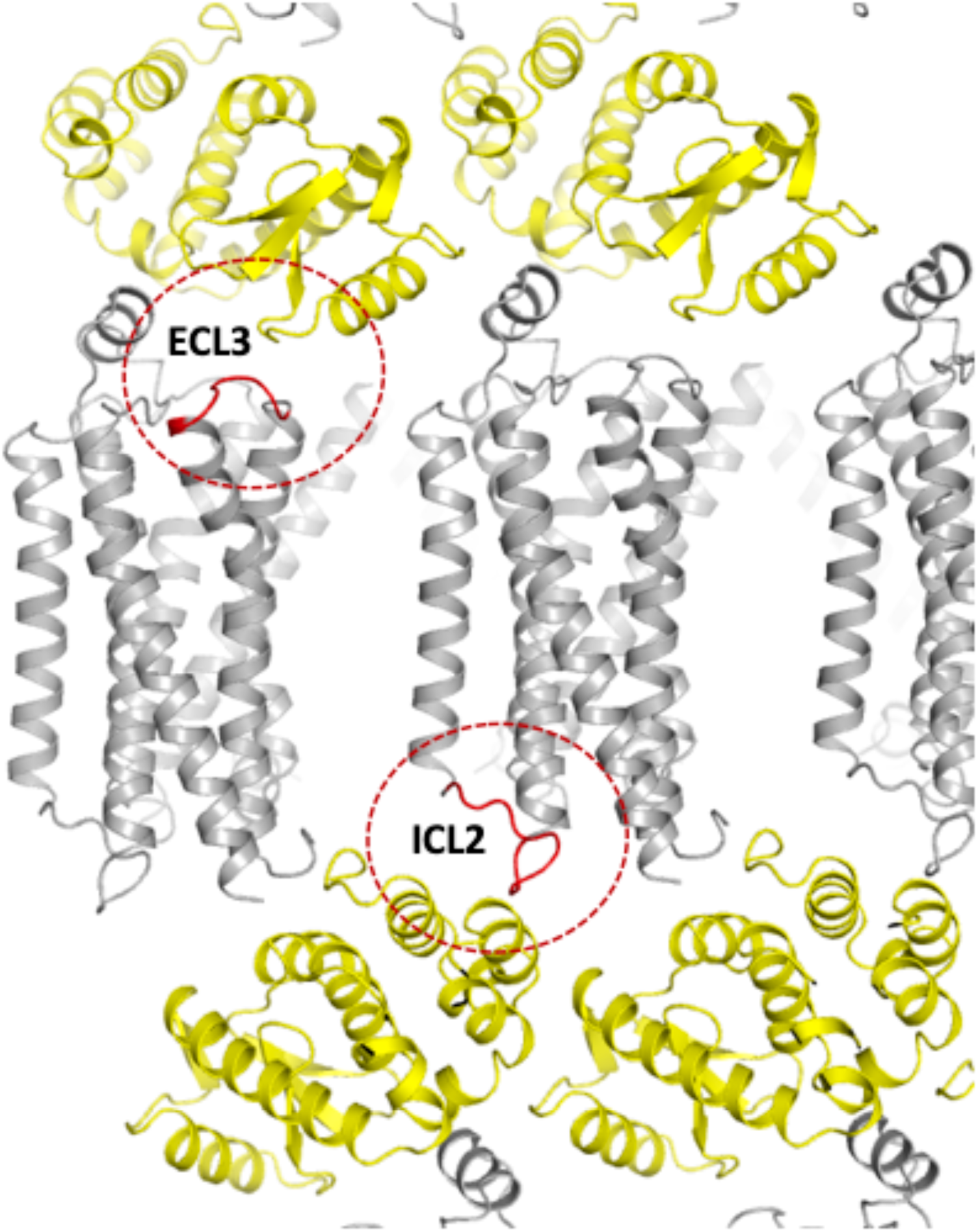
Crystal packing of the inactive structure of β2AR determined by X-ray crystallography. The β2AR is shown in gray and the T4L is shown in yellow. ECL3 and ICL2 are shown in red and highlighted with red circles.

**Fig. S4.**
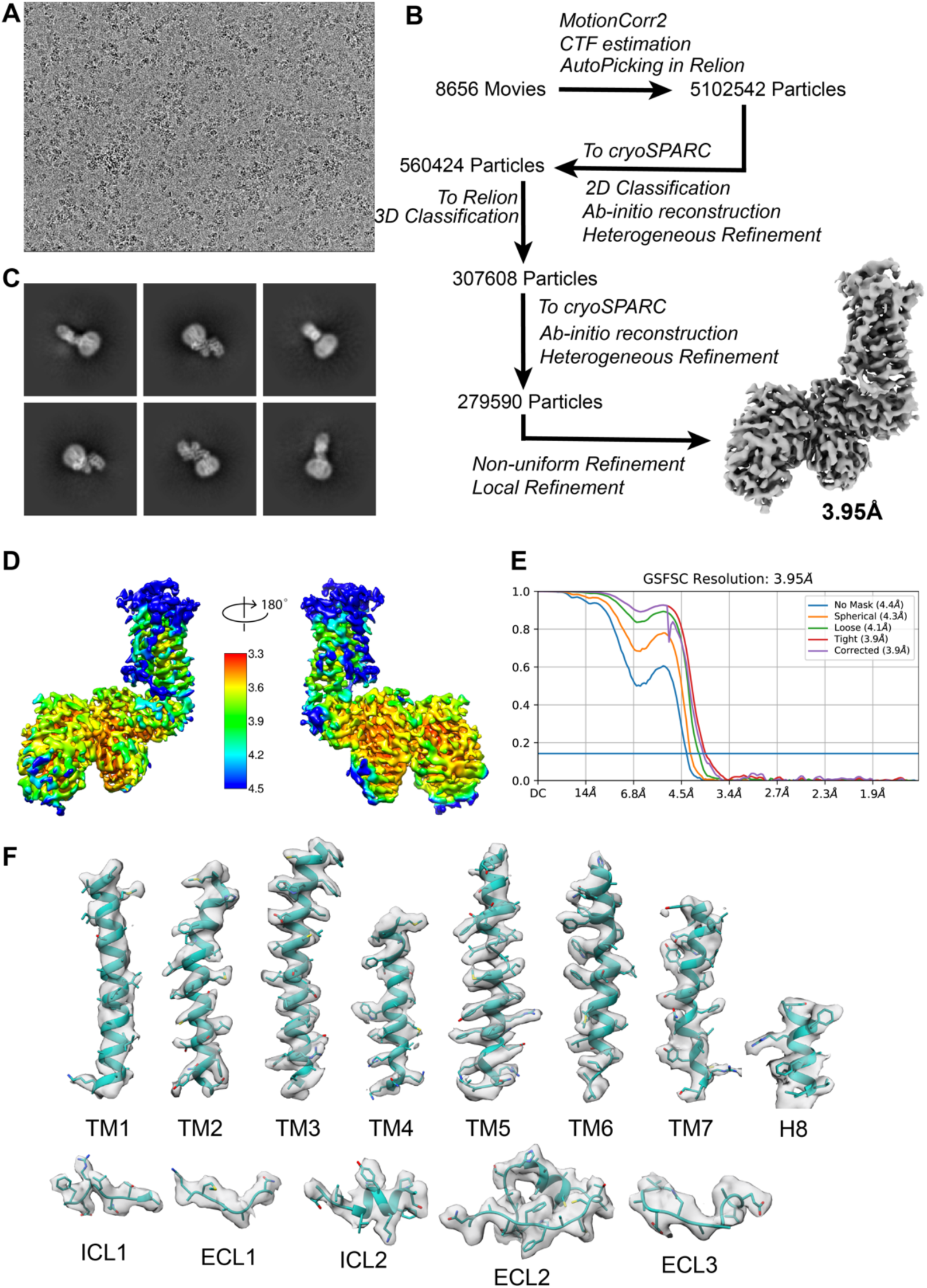
Cryo-EM data processing of apo-state β2AR-CN-FKBP12 complex. (A) Representative micrograph of the β2AR-CN particles. (B) Cryo-EM image processing workflow for image processing. (C) Representative 2D classification result. (D) Local resolution map viewed from two directions. (E) Fourier shell correlation (FSC) curve with the estimated resolution according to the gold standard. (F) Representative density maps and models for TM1-7, H8 and loop regions for apo-β2AR2AR.

**Fig. S5.**
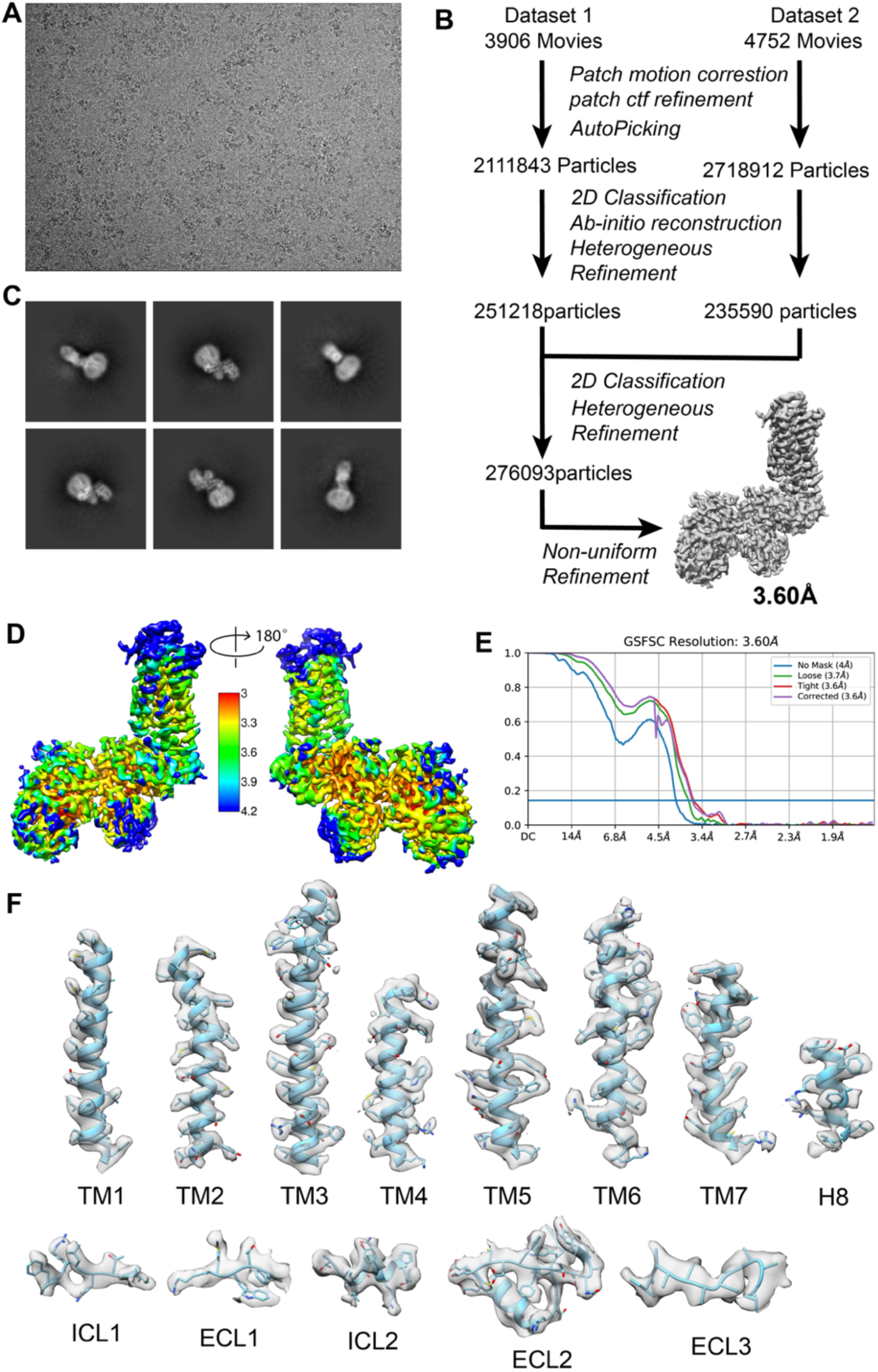
Cryo-EM data processing of norepinephrine-bound β2AR-CN-FKBP12 complex. (A) Representative micrograph of the norepinephrine-bound β2AR-CN-FKBP12. (B) Cryo-EM image processing workflow for image processing. (C) Representative 2D classification result. (D) Local resolution map viewed from two directions. (E) Fourier shell correlation (FSC) curve with the estimated resolution according to the gold standard. (F) Representative density maps and models for TM1-7, H8 and loop regions for norepinephrine-bound β2AR.

